# Snipers under stress: mentally simulated motor actions are resistant to acute stress in police officers

**DOI:** 10.64898/2026.01.04.697049

**Authors:** Hélène Monier, Sidney Grosprêtre, Nicolas Gueugneau

**Affiliations:** CEREN EA 7477, Burgundy School of Business, Dijon, France; Laboratoire C3S, Culture Sport Santé et Société, UFR STAPS Besançon, Université Marie et Louis Pasteur, Besançon, France; Institut Universitaire de France (IUF), Paris, France

**Keywords:** Police, acute stress, motor imagery, handgun manipulation

## Abstract

Motor imagery (MI) is a sensorimotor process allowing to mentally simulate a motor action and is widely used to enhance performance in domains such as rehabilitation, sport, and professional training. Although MI is increasingly incorporated into stress-management interventions, the reciprocal relationship, that is, the effect of acute stress on MI remains paradoxically poorly understood, particularly in ecologically valid settings. The present study investigated this issue in professional police officers performing physically and mentally a precise handgun manipulation task under graded stress conditions (psychological stress). Fourteen participants (mean age = 36.57 ± 2.10 years) was exposed to controlled, work-related stress scenarios designed to closely reflect real operational demands, while stress markers and temporal features of executed and imagined movements were recorded. Our results clearly indicate that motor imagery was preserved despite significant increases in stress levels, as the temporal characteristics of both executed and imagined movements remained stable across experimental conditions. These findings indicate that MI is resilient to acute psychosocial stress and support its relevance as a reliable training and performance-optimization tool in high-demand occupational settings.

## Introduction

Motor imagery (MI), the cognitive ability to mentally simulate movements without overt execution, plays a critical role in skill acquisition and motor performance [1–3]. In domains requiring fine motor control and high-precision tasks, the practice of MI can facilitate sensorimotor learning [4, 5], decision-making [6], and improve overall performance. Maximal voluntary force also increases after MI training [7]. At the neurophysiological level, many studies indicate that MI engages neural networks overlapping with those recruited in movement execution, including the supplementary motor area, premotor and primary motor cortices, and cerebellum - see Hétu et al. (2013) for a review. Learning a motor task by means of MI also prompts neural plasticity mechanisms within the motor cortex [9].

Despite its practical utility, the effectiveness of MI is contingent upon psychological and physiological factors. Mental and physical fatigue [10, 11], local pain [12], or circadian rhythms [12], can for instance alter MI and thereby reduce its effectiveness [14, 15]. High cognitive load and environmental stressors, such as external distraction or competitive pressure, can further interfere with MI by reducing attentional resources [16, 17]. Acute stress is characterized by transient psychophysiological responses to external stimuli. It is a state of homeostasis threat that occurs when environmental demands surpass the organism’s adaptive capacity, triggering behavioral adjustments to preserve the balance between autonomic branches [18]. From a neurophysiological perspective, a stressful stimulus activates both the hypothalamic-pituitary-adrenal axis and the autonomic nervous system (ANS); the sympathetic response increases, leading to an increase in cortisol and adrenaline levels, heart rate frequency, and arterial pressure [19, 20]. This, in turn, alters the activity of supraspinal structures, including the hippocampus, amygdala, and prefrontal cortex [20]; and ultimately shape sensorimotor performance. For instance, acute stress can increase muscle co-contraction [21], alter movement kinematics [22], and modify overall motor strategy [23], prioritizing rapid movements at the expense of accuracy [24]. These alterations are underpinned by cognitive mechanisms related to attentional control and working memory, which are compromised under acute stress. Attentional narrowing selectively enhances focus on threat-related stimuli [25], thus reducing peripheral awareness and impairing sensorimotor integration [26]. Simultaneously, stress-induced dysregulation of the prefrontal cortex could disrupts working memory, diminishing the brain’s capacity to adjust motor plans [27].

The relationship between stress and MI has attracted growing interest in the literature, particularly with regard to the potential of MI-based interventions as an effective strategy for stress regulation and performance optimization under pressure [28–30]. For instance, Page et al. (2016) demonstrated that a brief mental-skills training program incorporating MI significantly improved both memory recall and operational performance in police cadets undergoing high-stress training scenarios, relative to a control group. It is noteworthy that, despite these prior research critically examining the potential influence of MI on stress regulation, the reciprocal relationship has received limited attention, particularly within ecologically valid, work-related contexts. Actually, to the best of our knowledge only two experiments have explored the impact of acute stress on mentally simulated actions; one pilot study [31] and one randomized controlled trial [32]. While both works did not report significant effect of acute stress on MI capacities, these findings was based, however, on artificially induced stress within laboratory settings. Precisely, stress conditions were generated by immersing the hand in cold water (Cold Pressor Test), by answering arithmetical exercises while being filmed and judged by an audience (Trier Social Stress Test), or by the combination of both (Maastricht Acute Stress Test) [31–32]. Although these manipulations can prompt typical modulations of stress biomarkers (e.g. heart rate) [33], they are far removed from field situations, whether in sports competition, rehabilitation, or high-risk professional contexts. Moreover, in the previous studies [31–32] the temporal distance between the stressors and MI evaluation (i.e. > 20 min) makes it difficult to generalize the results to real-life situations, where it is appropriate to rapidly act, or plan to act under conditions of acute stress.

Some professional contexts, such as the police force, are particularly exposed to such demands. Police work and training, and particularly that of precision shooters (or snipers), provides an ideal opportunity to assess the effect of stress on MI in work-related contexts. Snipers can operate under extreme conditions of psychological and physiological stress [34]. Objective stressors include high-stakes decision-making (e.g. high-consequence decisions, determining whether to use lethal force under ambiguous conditions) [24, 35], emergency management (e.g. precise actions with short deliberation time) [36], or managing events endangering the agent’s integrity or life [37]. Social and organizational factors such as the expectation to perform flawlessly, combined with scrutiny from superiors or the public [38], and frequent shift work activities also contribute to the psychophysiological stress of those professionals [39]. It is now widely acknowledged that these factors can contribute to the long-term deterioration of both the physical and mental health of officers, underscoring the importance of integrating stress management strategies into police training programs [40]. Consequently, given the reliance of sniper preparation on MI-based rehearsal [28, 41], understanding the precise impact of acute stress on MI in work-related scenarios now becomes essential for optimizing training methodologies and operational readiness.

In this context, the present study aims to evaluate the impact of acute stress on MI performance in trained precision shooters. Using a controlled experimental design, we induced graded acute stress through a work-related context while assessing MI efficacy by means of behavioral measures, i.e. temporal congruency between executed and imagined motor actions. Psychophysiological indicators and psychometric scales were employed to quantify stress responses and MI vividness. Spurred by the few preliminary data on the subject, we hypothesized that there will be limited effect of acute stress on MI abilities.

## Material and methods

### Participants

Fourteen healthy right-handed male participants, aged between 24 and 50 years (36.57 ± 2.10 years) gave their written informed consent to take part in the experiment. The minimal sample size required to detect a significant stress-related difference of 15 beats per minute (± 10 bpm) in heart rate is equal to 6, with a level of α = 0.05 and power of 0.8 (Biostatgv software). This was determined based on previous published data focusing on performing behavioral tasks under stress [30]. Consequently, the current final sample size of n=14 ensured a reliable detection of variations in the selected stress biomarkers. Participants’ characteristics are summarized in table 1. All were experienced police officers with at least 3 years of activity and were recruited during the organization of a sniper training camp based in France. They all had a regular sport/physical activity (table 1). None of the participants reported suffering from sleep disorders, and their general sleep quality was self-estimated as moderate to good. Three participants were occasionally involved in night work for their operational activities, i.e. 1-3 events per month, but at the time of the study all reported having normal circadian rhythms with a diurnal activity and a nocturnal sleep. Drug intake (including alcohol, coffee, and tobacco) that could have altered mental or motor performance where prohibited 3 hours before the experimental sessions. Participants had no history of neurological disease, and did not have severe musculoskeletal injury within the last year. All participants received a detailed explanation of the study and informative note before the experiment, and provided a written consent form in agreement with the Declaration of Helsinki [42]. It should be noted, however, that details concerning the stress induction were kept confidential until the very end of the protocol in order to guarantee the potential effects of stress on the dependent variables [43]. The ethics of this research have been approved by the operational direction of the Police Company CRS 40, (*Compagnie Républicaine de Sécurité*, Plombières-lès-Dijon, France).

**Table 1.** Sample characteristics of the participants. *Note*. Values expressed as mean (± SD). Abbreviations: STAI-Y-B, State-Trait Anxiety Inventory *form* Y *part* B (trait). MIQR, Movement Imagery Questionnaire Revisited.

| General features |  |
| --- | --- |
| Age (years) | 36.57 ± 2.10 |
| Seniority in Police (years) | 10.07 ± 1.98 |
| Physical activity (hours per week) | 7.25 ± 1.15 |
| Sleep (hours per day) | 7.03 ± 0.79 |
| Anxiety |  |
| STAI-Y-B | 36.93 ± 6.14 |
| Motor imagery ability |  |
| MIQ-R *Global* | 44.37 ± 4.86 |
| MIQ-R *kinesthetic* | 21.42 ± 3.97 |
| MIQ-R *visual* | 23.21 ± 4.12 |

### General experimental design

This research was part of a training module to select snipers from the French National Police; i.e. the GPIS4 module (4th Generation Protection and Intervention Section) which is a 3 weeks training program including physical and operational tests related to precision shooting. The experiment had four sessions involving psychosocial and sensorimotor evaluations. The sessions lasted a maximum of 15 minutes. An interval of at least three days separated the three experimental sessions, and the order between control and stress sessions was pseudo-randomized (7 participants did the control first, and 7 did it last).The first session included preliminary assessments while the three others specifically evaluated MI in stress-varying conditions. Preliminary assessments were done individually and were supervised by one of the authors, at least one week before the other three experimental sessions. The three experimental sessions took place at participant’s workplace, where stress level was experimentally modulated. Then, participants underwent a control session, a low-stress session, and a high-stress session (see details hereafter). All experimental sessions were scheduled between 10 a.m. and 6 p.m. to control potential circadian effects [13, 15, 44]. Physical exercise and important cognitive solicitations were proscribed before the testing sessions, i.e. at least 1h. Figure 1A illustrates the general timeline of the experimental design.

**Figure 1.**
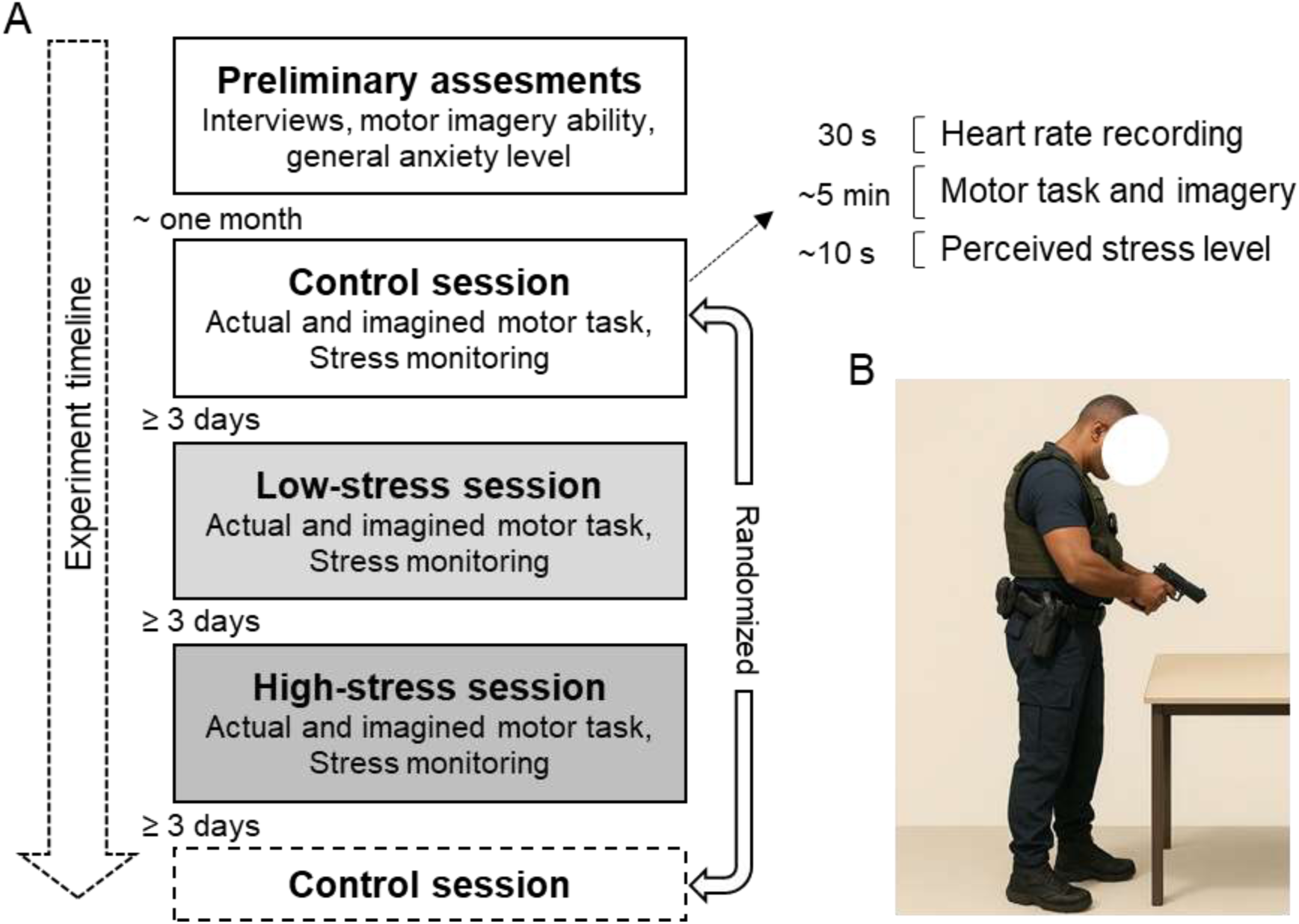
Experimental design. **A**. Timeline of the experimental sessions (left), and sequence of the behavioral and physiological measures (up right). **B**. Illustration of a participant executing the handgun manipulation task (computer-generated and royalty-free picture).

### Preliminary assessments

Preliminary assessments included i) formal interviews, ii) a general evaluation of stress level, and iii) subjective evaluations of MI capacities. Formal interviews enabled to collect elementary demographic data, general sleep quality, weekly physical activity level, recent medical history, and years of seniority in the police. General stress level was assessed with the trait anxiety questionnaire STAI-Y-B [45]. The STAI-Y-B evaluates a global level of stress as perceived in daily life, according to twenty questions of 4-items. The score ranges from 20 to 80 and reflects a self-reported state of habitual stress and allow detecting potential anxious profiles or depressive state among participants, e.g. those who obtain a score ≥56 [45]. MI ability was measured by the Revised Movement Imagery Questionnaire MIQ-R [46]. During MIQ-R, the participants performed physically and mentally body movements, and had to use either visual MI (i.e., third person perspective) or kinesthetic MI (i.e., first person perspective with body sensations usually associated with the actual execution of the action). Participants rated the quality of their mental representation with a 7-point scale in 8 items (maximum score = 56).

### Experimental sessions

The core experiment consisted in evaluating the ability to form explicit mental motor images during a professional scenario while varying stress levels. A well-known mental chronometry paradigm [11, 13, 15, 16] was used to compare actual and imagined movement durations during a standardized professional motor task (see *Motor task* hereafter). Three distinct contexts allowing to modulate the level of perceived stress were used (see *Stress induction* hereafter). These contexts thus constituted three separate experimental sessions.

#### Motor task

The task consisted in executing a *commissioning* of the service handgun (Sig-Sauer SP 2022); Figure 1B. This task refers to a precise movement sequence involving strictly codified motor actions. Basically, it involves the following key steps that are realized sequentially from standing upright in a neutral anatomical position: 1. grab the weapon from the belt, 2. operate the breech twice checking that the barrel chamber is empty, 3. control visually the general functioning of the weapon (hammer, breech retainer, magazine), 4. load, and operate the breech, 5. disarm the hammer by lowering the disarming lever, 6. control visually and by touch the loading indicator, 7. replace the weapon to the belt.

This motor task was selected because it met two important criteria from ergonomics and neuroscientific points of view. First, it is a well-defined professional gesture of the police forces; and second, the task is rehearsed regularly during the career of police officers (∼300 times per year; that is a mean of ∼3000 times at a group level for the participants). It implies that its execution is well integrated within the motor repertoire of the participants – i.e. based on accurate motor representations and poorly sensitive to learning processes in the present experiment. This ensured a negligible learning effect across the trials and the different experimental sessions, of which the order was, in any case, randomized.

For the actual movements, participants had to perform the entire movement sequence at a natural pace, as they would do in a professional context. For the imagined movements, they were instructed to feel themselves (kinaesthesia) performing the motor task (first person perspective or internal imagery) rather than just watching themselves executing it (external imagery). After 1 to 2 trials of familiarization, all participants declared being able to generate mental motor images. As during actual trials, to provide congruent visual information across conditions participants kept their eyes open during MI and had to fix their gaze on a point on the table in front of them. Importantly, experimental studies provided evidence that there was no direct influence on motor performance relative to the open eyes’ condition [48]. To note, closing eyes during the imagery task does not prevent visual tracking, and abilities to perform efficient MI are not dependent of eyes movements [46]. The participants were requested to realize eight actual trials and eight imagined trials in a random order in such a way that all 16 trials were mixed together. During both actual and imagined trials, participants were requested to say “Go” when they started to move, and “Stop” when they finished the whole movement sequence. The sequence of trials was arranged according to the participants’ preferences, and breaks were included as needed. The experimenter recorded sequence durations from the “Go” to the “Stop” signals using a digital stopwath (resolution 1/100 s).

#### Stress induction

Stress was induced by controlling the environment in which the motor task was realized (i.e. control session, low-stress session, and high-stress session). In the control session, the participants realized the motor task in a standard classroom of the Police training center. The environment was calm, and the ambient temperature was ∼20°C (for the whole sessions). They were only accompanied by the main experimenter who adopted a neutral behavior, characterized by a standardized tone of voice, absence of evaluative feedback, and emotionally non-expressive facial and postural cues, in order to avoid influencing participants’ affective or stress responses. The location of the session, and the way the experimenter interact with participants constituted a neutral context. Apart from the instructions concerning the execution of the motor task (see chapter *motor task*), no other indication was given (e.g. reward, punishment).

In the low-stress session, the participants performed the motor task during the marksmanship training camp in the presence of the main experimenter and the chief of the training operations. The session was done in a large meeting room before the participant had to practice physical or shooting exercises. The chief attended the entire session from the participant sidelines. He intervened only at the beginning of the session by explaining that the motor task had to be carried out with caution, as it was an important part of the operational training.

The high-stress session implied the same procedure as the low-stress session with the crucial difference that at the beginning of the session the chief indicated that any error during the gun manipulation would lead the participant to be excluded from the rest of the training camp. The chief adopted a predetermined distant and cold behavior. Such authoritative behavior was important to reinforce the social aspect of the stress induction [43]. Also, the threat of exclusion from the camp was expected to significantly increase the level of stress, as this event constitutes an important achievement in the career of the participants; i.e. basically, such a training camp is carefully prepared for months and can allow the policemen, if the issue is positive, to progress significantly in terms of responsibility, expertise, and career. Importantly, the threat of exclusion from the camp was not real; this information was revealed to the participants at the end of the experiment. Before each experimental session, the participants were required to remain seated for two minutes while receiving instructions, in order to standardize their basic physiological state.

#### Monitoring of stress induction

cardiac activity and perceived stress level were used as stress markers. Cardiac activity was indexed from the heart rate (bpm) using a smartwatch equipped with a cardiofrequencimeter (Garmin®, Forerunner 45). The average heart rate over 30 s preceding the first trial of the motor task was calculated. Perceived stress level was recorded by using a visual analog scale (VAS) to report how stressful the participant experienced the situation (from 0 “absolutely not stressed” to 10 “extremely stressed”). VAS value was recorded immediately after the last trial of the motor task (see Figure 1A for the precise timeline).

### Data analyses

*Statistics* -Statistics were performed using the software *JASP* (0.19.3). Mean values at the group level ± standard deviations (SD) were computed for the general demographic parameters, anxiety trait scores from the STAI-Y-B, and MI ability scores from the MIQ-R. For each participant, average values (± SD) were computed in each condition for heart rate, perceived stress level, and the durations of executed and imagined movements. An Index of Motor Imagery Performance (IMIP), calculated by dividing the durations of imagined movements (MD_Ima_) by the durations of executed movements (MD_Exe_): IMIP = MD_Ima_ / MD_Exe_, allowed to evaluate the temporal equivalence between executed and imagined movements across experimental conditions. Thus, an IMIP equal to one indicates a perfect temporal equivalence between both movement modalities, while an IMIP equal to 0.95 indicate that the duration of the imagined movement is 5% shorter than the one of actual movement. Correlation analyses between executed and imagined durations were also done by entering all trials in Pearson’s analyses. The temporal variability of executed and imagined movements was assessed by means of a Coefficient of Variation for each experimental condition (CV; defined as the standard deviation divided by the average duration, multiplied by 100). The normality of the distribution of the variables was verified by means of a Kolmogorov–Smirnoff test and by visually inspecting the Q-Q plots of the model residuals; sphericity was verified by a Mauchly’s test.

To consider the effect of *stress* (control, low-stress, high-stress) on the stress score and heart rate, a 1-way repeated measures analysis of variance (ANOVA) was performed. To consider the effect of *stress* (control, low-stress, high-stress) and movement *modality* (executed, imagined) on movement duration and CV, a 2-way ANOVAs was performed. Finally, a 1-way (stress effect) ANOVA was performed on the IMIP. These analyses were done on the averaged group values (n = 14). For all the analyses, post hoc differences were assessed by means of Bonferroni tests. Significance was set at P < 0.05. Correlation analyses between the whole executed and imagined durations were also done. The effect size was evaluated by calculating the η_p_² (partial eta squared, with thresholds of 0.01, 0.06, and 0.14 for small, medium, and large effects) for the ANOVA analyses, and the Cohen’s *d* for post hoc tests (with thresholds of 0.20, 0.50, and 0.80 for small, medium, and large effects).

## Results

### General features

Table 1 summarizes the demographic and general characteristics of the participants. The mean STAI-Y-B score indicated normal trait anxiety levels (as compared to the general population [45, 49]), with no participant scoring ≥ 56 (threshold for elevated anxiety). In addition, the mean MIQ-R score reflected good motor imagery abilities [44]; see Table 1.

### Stress markers

Figure 2A and Figure 2B depict, respectively, the perceived stress level, indexed by the stress score, and heart rate across the three experimental sessions. Both measures exhibited a marked increase from the control session to the high-stress session. Repeated-measures ANOVA confirmed a significant main effect of *stress* on both the stress score (*F*(2, 26) = 25.45, *p* < 0.001, η_p_² = 0.66) and heart rate (*F*(2, 26) = 21.73, *p* < 0.001, η_p_² = 0.62). Post-hoc comparisons indicated that stress scores and heart rates were significantly higher during the high-stress session compared with the low-stress session (*p* < 0.01, *d* ≥ 1.25 for each comparison) and the control session (*p* < 0.001, *d* ≥ 2.16 for each comparison). Both indices were also significantly higher in the low-stress session relative to the control session (*p* < .05, *d* ≥ 0.91 for each comparison).

**Figure 2.**
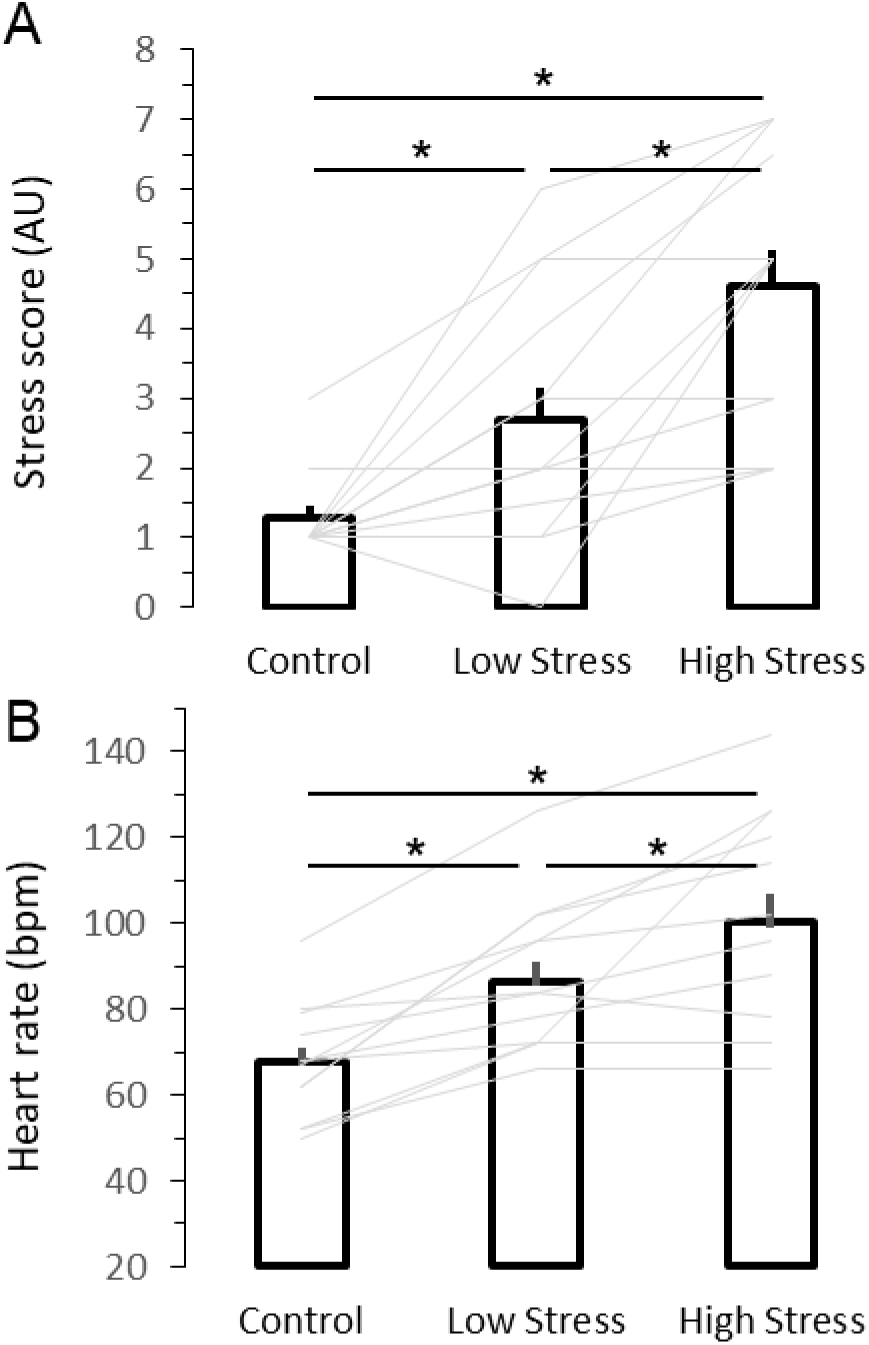
Stress markers. **A**. Mean values (± SD) and individual data (histograms and grey lines, respectively) of the stress score are represented as a function of the stress sessions. **B**. Mean values (± SD) and individual data (histograms and grey lines, respectively) of heart rate are represented as a function of the stress sessions. * indicates a significant difference with p < 0.05.

### Temporal features of executed and imagined movements

Table 2 presents the mean durations of executed and imagined movements during each of the three sessions. A two-way ANOVA revealed a significant main effect of *modality*, F(1, 13) = 66.35, p < 0.001, η_p_² = 0.37; while neither the main effect of *stress*, F(2, 26) = 0.51, p = 0.95, η_p_² = 0.002, nor the interaction effect, F(2, 26) = 0.33, p = .69, η_p_² = .002, reached significance. Overall, the duration of executed was longer than those of imagined movements. The relationship between executed and imagined movement durations is further illustrated in Figure 3. Temporal equivalence between the two modalities (as expressed by the IMIP, Fig. 3A) remained stable across the three sessions, averaging 0.83 ± 0.10, as confirmed by a one-way ANOVA, F(2, 26) = 0.43, p = 0.71, η_p_² = 0.026. Moreover, correlation analyses (Fig. 3B) showed consistently high and significant correlations between executed and imagined durations across all the three sessions (*r* > 0.8; p < 0.01 for each correlation coefficient).

**Table 2.** Durations of executed and imagined movement as a function of stress. *Note*. Values expressed as mean (± SD).

| Experimental sessions | Control | Low Stress | High Stress |
| --- | --- | --- | --- |
| Executed movement (s) | 10,06 ± 0,33 | 9,86 ± 0,39 | 9,82 ± 0,36 |
| Imagined movement (s) | 8,23 ± 0,43 | 8,15 ± 0,39 | 8,30 ± 0,50 |

**Figure 3.**
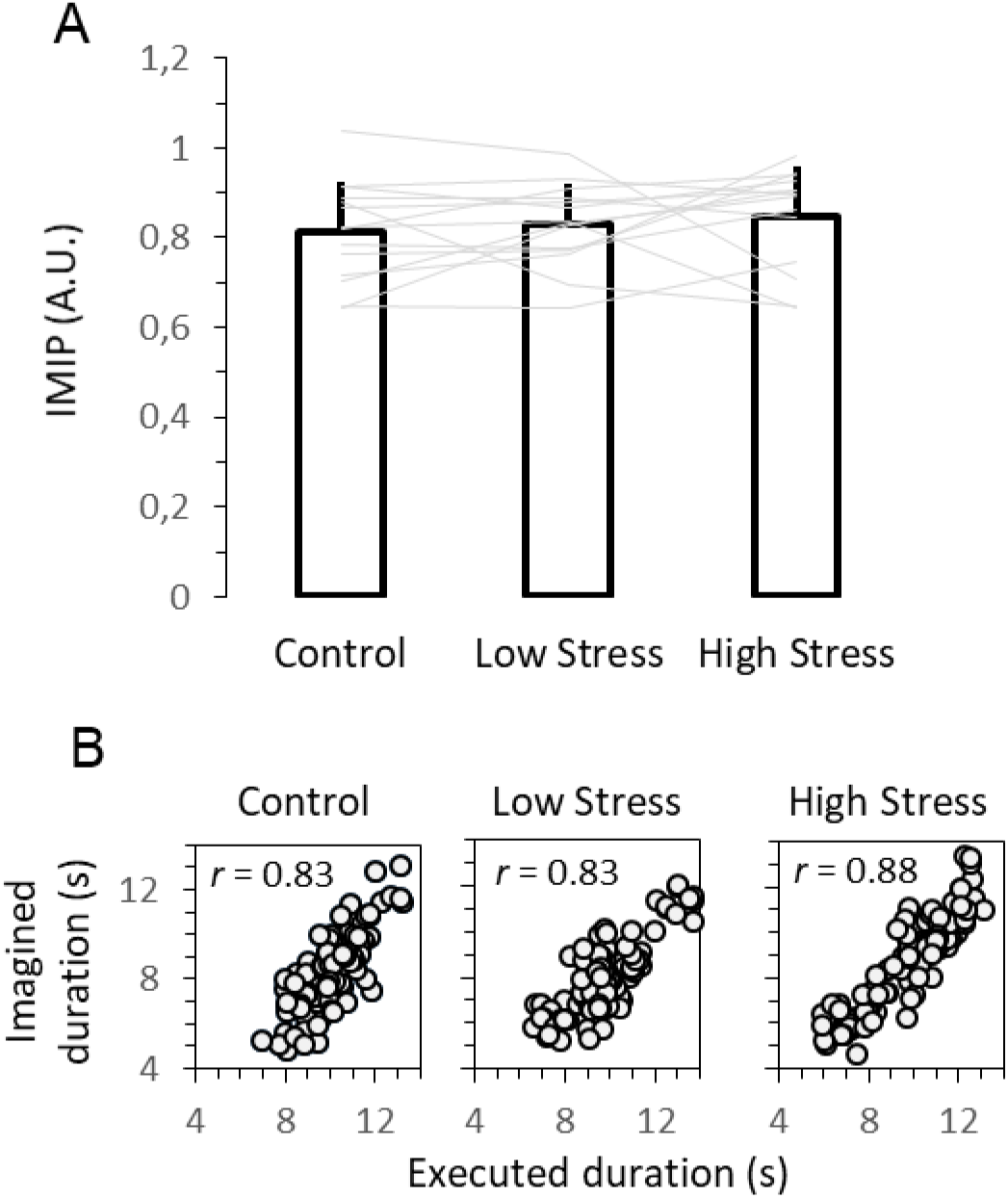
Motor imagery as a function of stress. **A**. Mean values (± SD) of IMIP and individual data (histograms and grey lines, respectively) are represented as a function of the stress sessions. **B**. Duration of each executed movement is plotted vs. duration of the corresponding imagined movement for all subject in each stress session. The correlation coefficients are reported nearby the corresponding scatter plot.

**Figure 4.**
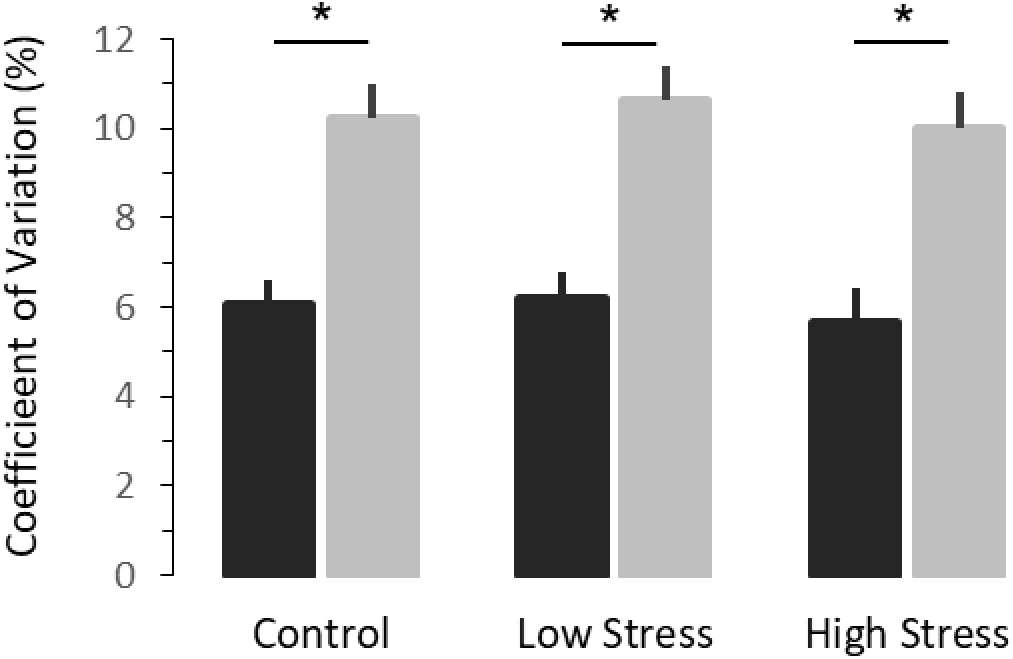
Temporal variability of executed and imagined movements. Mean values (± SD) of the coefficient of variation for executed and imagined movements (black and grey histograms, respectively) are represented as a function of the stress sessions. * indicates a significant difference with p < 0.05.

Figure 5 depicts the average values of the coefficient of variation (CV) during the three sessions. A two-way ANOVA showed a significant effect of *modality* (F(1,13) = 144.8, P < 0.001, η_p_² = 0.53), while neither the main effect of *stress*, F(2, 26) = 0.31, p = 0.73, ηp² = 0.007, nor the interaction effect, F(2, 26) = 0.042, p = 0.95, η_p_² = 0.001, reached significance. CV thus remained stable across sessions, and was significantly higher for the imagined (on average, 10.28 ± 0.77) than the executed movement durations (on average, 5.96 ± 0.64).

Finally, to evaluate potential learning or adaptation across trials, the mean durations from the first two (trials 1–2) was compared to the last two (trials 7–8) repetitions within each session and modality. No significant differences were observed (p > 0.10 for all comparisons), suggesting an absence of systematic learning effect.

## Discussion

The present study aimed to assess the effect of acute stress on MI in precision shooters (or snipers). To this end, the temporal features of executed and imagined movements of a handgun manipulation task were evaluated across three experimental sessions, involving ecological scenarios with gradual stress intensities. The main findings pointed out that while the stress manipulation was effective, as evidenced by significant increases in both subjective stress ratings and heart rate, the temporal correspondence between executed and imagined movement durations remained stable across stress sessions. Also, the duration, and the temporal variability of movements, whether executed or imagined, appeared to be independent of the stress level. Together, these results suggest that while acute stress reliably impacts physiological and psychological stress markers, it does not compromise the temporal congruence of MI in this population. The data confirmed previous laboratory-based studies in healthy adults and further highlight the potential and the feasibility of MI in stressful professional contexts.

The robustness of the stress induction procedure was confirmed by convergent evidence from subjective and physiological measures. Indeed, both perceived stress and heart rate showed gradual and significant increases across the three experimental sessions. These findings align with well-known stress-induction paradigms such as the Trier Social Stress Test (TSST), the Socially Evaluated Cold Pressor Test (SECPT), or the Maastricht Acute Stress Test (MAST), which reliably elicit both psychological strain and cardiovascular activation [31–33, 50]. Importantly, the present setup provides substantial advantages over the two prior studies that examined MI under stress conditions [31, 32]. First, a graded stress induction with low- and high-stress sessions was used, while the previous experiments employed binary “all or nothing” stress paradigms, which may have prevented the detection of a threshold effect (i.e. depending on stress intensity). For instance, during the present high-stress session heart and perceived stress level increased by ∼40% and ∼400% respectively, as compared to the control session; whereas Schlatter et al. (2020) reported more modest increases of ∼20% and ∼10% for the same variables measured concurrently. Second, the main dependent variables used here, i.e. the executed and imagined movements’ durations, were assessed right after the stressor, while the previous studies included a ∼15-20 min time interval between the stressor and task evaluation [31, 32]. Such a delay allows the long-lasting activation of the hypothalamic-pituitary-adrenal axis, with a peak in cortisol levels ∼20-30 min after the stressor; yet, it may mask faster responses related to the activation of the sympathetic adreno-medullary system, with an immediate rise in cardiovascular activity and a return to baseline after ∼15-30 min [19, 33, 50]. By doing so, it was ensured that executed and imagined movements were completed during the concurrent activation of the ANS and hypothalamic-pituitary-adrenal axis, i.e. right after the stressor. This point has been raised in several studies which have shown that acute stress alters cognitive activities (e.g. working memory) when the task is evaluated right after the stressor [51, 52], but not after a 20 min time interval [53, 54]. Trapero-Asenjo et al. (2024) also highlighted that the preliminary results linking stress and MI needed to be evaluated in a context where the task is assessed right after the stressor.

The present main findings confirm that acute stress did not alter the temporal relationship between executed and imagined movements. In every sessions, the durations of imagined movements remained closely aligned with the durations of executed movements, yielding high correlations and stable Indices of Motor Imagery Performance (IMIP). These results confirmed the hypothesis and are in line with preliminary studies reporting similar data in other populations [31, 32]. These data can hardly be attributed to insufficient stress level (as mentioned above), or to a lack in task complexity. Indeed, the handgun manipulation task involved precise series of fine and complex hand and fingers movements; while the previous works used simple arm pointing movements [31] or basic tasks from the motor repertoire (e.g. jumping, flexion and extension of the knee and hip) [32]. The apparent resistance of MI to acute stress contrasts with a large body of evidence showing that stress impairs numbers of higher-order cognitive functions [55]. This discrepancy may originate from the distinct neurophysiological substrates of MI, which predominantly relies on cortical sensorimotor networks [2, 8]. Acute stress has been shown to mostly disrupt the functions of the prefrontal cortex and hippocampus [20, 27, 56], thereby compromising executive processing -including working memory, attention, cognitive flexibility, decision-making [55] - and visuospatial processing [57], respectively. By contrast, MI recruits neural networks that overlap with motor execution, including premotor, parietal, and subcortical regions such as the basal ganglia and cerebellum [8]. These neurophysiological distinctions may therefore account for the apparent resilience of the sensorimotor processes at work during MI to transient stress-related impairments. Interestingly, however, mental rotation tasks, regulated by *implicit* MI processes [58], are impaired by acute stress [31]. Such tasks specifically require important visuospatial processing and rely on hippocampal engagement [59], whose function is known to be compromised under stress [56].

Moreover, the temporal features of executed and imagined movement gave two remarkable results. First, imagined movements consistently exhibited greater variability than executed movements, as quantified by CV. This observation replicates prior research showing that MI is intrinsically more variable than overt execution [5, 15, 44, 47]. As discussed in our earlier works, this increased variability can be accounted for within the framework of motor prediction [60, 61]. During actual execution, the central motor command predictively sets the spatiotemporal structure of voluntary movement but also benefits from sensory feedbacks, which allows continuous regulation of movement’s dynamics. In contrast, MI relies exclusively on sensorimotor predictions generated by forward models, which are neural networks that simulate the dynamics of the body and its interaction with the environment [60]. Because no actual movement occurs during MI, the efference copy-based predictions cannot be updated through sensory feedback. Consequently, trial-to-trial variations in imagined movement remain uncorrected, resulting in the high temporal variability observed during MI. Second, the duration and temporal variability of both executed and imagined movements remained stable across the experimental sessions. These data partly contrast with studies in which sensorimotor performances deteriorate with stress. For instance, fine motor control tasks (e.g. surgery acts [62], precise force control [63], or handgun shooting [24, 35]), and gross motor skills (e.g. locomotion [64], throwing tasks [65]), have been shown to exhibit significant impairments under acute stress. Yet, such tasks rely heavily on visuomotor attentional mechanisms and continuous sensory adjustments. These ongoing and dynamic integration of sensory inputs is known to be particularly disrupted by stress [66]. Conversely, the handgun manipulation task used in this study is a highly practiced skill, as confirmed notably by the absence of a learning effect and the trial-by-trial stability of both executed and imagined movements. The task may thus likely rely on *automatic* sensorimotor loops rather than stress-sensitive prefrontal control processes. Over learning and consolidation, the control of such motor sequences shifts from prefrontal and attentional networks towards primarily subcortical loops involving the cerebellum and basal ganglia, supplemented by sensorimotor cortical representations. [67]. Also, the task, being performed at a natural and comfortable pace, probably imposed low demands on attention and working memory [68]. Generally, by showing that the temporal congruence between actual and imagined actions remains stable under acute stress, and that the latter did not exacerbate the variability of both movements, our findings further suggest that stress may spare sensorimotor circuits involved in well-learned motor routines.

An alternative standpoint, coming from the literature in psychology, may relate to population-specific coping abilities in managing stress. Coping refers to cognitive and behavioral adjustments aimed at managing perceived threats and may operate deliberately or automatically [69]. Police officers, who are repeatedly exposed to high-risk situations, typically develop strong coping skills and preferentially rely on active, task-focused strategies rather than emotion-focused or avoidance coping, which is associated with poorer performance [70, 71]. Large-scale studies in police populations have shown a predominance of adaptive coping strategies under stress [72, 73]. Although cultural differences may exist, such coping profiles appear consistent across law enforcement contexts [71]. Thus, in the present study, the career-related nature of the stress induction may have further reinforced task-focused coping, contributing to the absence of stress-related changes in task performance.

Our study has, however, some limitations. Electromyographic measurements could have been done, particularly to control for the absence of muscle contractions during MI. Yet, implementing such an evaluation could have significantly increased the duration of each session, and thus undermined our ecological approach. Also, more sensitive stress markers such as heart rate variability (e.g., RR intervals, RMSSD), or galvanic skin response, could have been used. Nevertheless, reliable RMSSD estimation requires longer recording periods (≥5 min), which exceeds the analysis windows used here [74]. In contrast, galvanic skin response provides a valuable index of stress but is difficult to disentangle from physical effort and is best recorded at rest. Cortisol levels may also serve as a robust physiological marker of stress. Future investigations could also address whether the task employed was too *easy* by introducing more complex sequences, or those for which the subjects are naive.

To conclude, our study confirm that MI is preserved under acute stress. Notably, the current paradigm was designed to mimic operationally relevant contexts for police officers, thereby increasing its ecological validity. Acute stress is an inherent feature of law-enforcement activities [34, 37, 39, 40], and validating experimental manipulations in this population represents an important step for translational applications [75, 76]. Our data thus support the use of MI techniques in training and operational contexts. Of note, our stress-induction methodology is closer to protocols involving *social* stress components – e.g. being seen and/or judged by peers or an audience; than protocols involving *physical* components – e.g. sensory alterations, physical effort or simulated danger. Future works should include physical stressors (e.g. fatigue, perceived danger).

## Data availability statement

The datasets generated and/or analyzed during the current study are available from the corresponding author upon reasonable request.

## Acknowledgments

We would like to thank the Training Center of Precision shooters (Plombières-les-Dijon, France) for opening their doors during in situ observations and experimentation. We would also like to thank our partner, the Research Laboratory of the French Police Academy (Ecole Nationale Supérieure de la Police, France), through whose agreement to host us at the Training Center formalized our visit. Hélène Monier was supported by the Region Bourgogne Franche-Comté (France) – “Projet TIGRE”, ANER. Nicolas Gueugneau was supported by the Region Bourgogne Franche-Comté (France) -”Amorçage-Envergure”, PARCK-CognAction (UB 905 -CR 0900).

## Author contributions

HM and NG designed and conduced the experiments. NG analyzed the data. HM, SG, and NG wrote the manuscript.

## Additional Information

The authors declare no competing interests.

